# A simple modification of library length for highly divergent gene capture

**DOI:** 10.1101/080689

**Authors:** Xing Chen, Gang Ni, Kai He, Zhao-Li Ding, Gui-Mei Li, Adeniyi C. Adeola, Robert W. Murphy, Wen-Zhi Wang, Ya-Ping Zhang

## Abstract

Hybridization capture is considered very cost- and time-effective method for enriching a massive amount of target loci distributed separately in a whole genome. However, divergent loci are difficult to enrich for the sequence mismatch between probes and target DNA. After analysis the distributional pattern of divergent loci in mitochondrial genomes (mitogenomes), we notice that the relatively variable regions are intercept by the relatively conservative regions. We propose to extend the length of library to overcome the problem. By using a home-made probe set to bait amphibian mitogeneomes DNA, we demonstrate that using 2 kb DNA libraries generate high sequence coverage in the highly variable regions than using 400 bp DNA libraries. These suggest that longer fragments in the library generally contain both relatively variable regions and relatively conservative regions. The divergent part DNA along with conservative part DNA is captured during hybridization. We present a protocol that allows users to overcome the gap problem for highly divergent gene capture.

## 1. Introduction

Target-sequence enrichment coupled with high-throughput sequence is very popular method in many disciplines, including evolutionary biology, population genetics, and human disease (1-3). Hybridization capture is considered very cost- and time-effective for enriching a massive amount of target loci distributed separately in a whole genome (4). There are exist versatile approaches to various target loci of interest, including exon-capture, ultra-conservative elements and home-made probe capture for any specific gene (2-3). Moreover, there is no need of *a priori* gene sequence of a species of interest. Currently, DNA sequence from closely relative species available in public databases allows researchers to design probe to bait target loci (5).

The general experimental steps of in-solution hybridization capture method is as follows: 1) prepare a DNA/RNA library from species of interest; 2) prepare probe to bait target genes; 3) then mix the probe and the library for enriching target genes; 4) sequence the enriched library. Sequence mismatch between probe and target sequence greatly influences gene coverage. Divergent gene region is difficult to design suitable probes referred to the closely relative species (5). Some studies report various ways to overcome the problem, such as optimize hybridization temperature (6-8), re-capture strategy (9), and probe optimization (10). Various temperature, including standard hybridization temperature 65 °C, low temperature 60 °C, 50 °C, 48 °C and 45 °C, touchdown approach are tried (1, 6-8). But temperature has been proved not the crucial parameter (5). In a mitochondrial genome research (1), we notice that gene conservativeness has striking relation with gene coverage. The most conserved regions, such *12s* and *16s rRNA*, have extremely high sequence-depth. In contrast, control region and NADH dehydrogenase genes have low or even no coverage. These suggest that sequence-depth is enough high for the mitogenome. The lack of read mapped to the divergent gene results from the extremely low efficiency of probe binding with the DNA fragments. Therefore, we focus on the molecular experiment solution instead of computational method to overcome the problem. To clearly understand divergent regions distributed in mitogenomes, we select 40 loci from 13 protein-coding genes and two rRNA genes in 33 mitogenomes from 24 amphibian genera. The length of variable regions are range from 400 bp to 1021 bp. The divergent regions are intercepted by the conservative region. The distributional pattern indicates that shearing the genome longer enable fragments generally contained both relatively conservative part and relatively divergent part.

In this study, to address the problem of gap in divergent genes, we hypothesis that extending library length to hybridization (termed LR-HY) is an effective optimization for improving the coverage of high divergent gene. In the standard protocol of in-solution capture hybridization, the length of the library is usually range from 200 to 500 bp for the convenient of the high-through sequence. This parameter is not suitable for the gene contained long divergent fragment. Just like the example of amphibian mitogenomes as above. To testify the hypothesis, we choose amphibian mitogenomes to illustrate the performance of LR-HY. The reasons we choose this animal group are: 1) the lengths of control region are varied from approximately 100 bp to more than 2kb; 2) the sequence of whole mitogenomes are more variable as compare to them from Aves and Mammalia; 3) Gene rearrangement of NADH dehydrogenase genes are common in many families. We compare the modified method with standard method with two species *Rana sp.1* and *Onychodactylus sp.1*. There are two gaps existed in the mitogenome of *Rana sp.1* and four gaps in *Onychodactylus sp.1* at these highly variable loci (Figure 2 A and B, orange line) when we use standard method. Instead, the LR-HY greatly improves the sequence coverage in these regions (Figure 2, green line).

The general pipeline of this study is shown in Figure 1A. A suitable probe set and long DNA library are necessary to prepare first. The probe set is randomly generated from Long-range PCR (LR-PCR) products since it is cost-effective and convenient to prepare. We designed a new set of universal primers by using mitogenomes from Amphibia, Aves and Mammalia. Two PCR reactions cover a complete mitogenome from any vertebrate species. To prepare long library, we shear a total DNA to the length of approximately 2-3 kb depended on the length of control region of amphibian mitogenome. Then the home-made probe is used to enrich target long DNA fragments in the library. In the downstream sequence experiment, an Ion Torrent Personal Genome Machine (PGM) is used to sequence because it is fast and relatively inexpensive in terms of each run (not price per base). Each run using 316 chip generated over 800 Mb for 60 samples and the data size for each sample is more than 10 Mb in general. These generated data are sufficient for *de novo* assembly of a complete mitochondrial genome. Although we sequenced with Ion torrent platform, the protocol could also be applied to Illumina platform with accordance with its library construction protocol. The detail LR-HY protocol is as follow. This protocol can be carried out in any molecular biology lab with standard library construction equipment.

**Figure 1.**
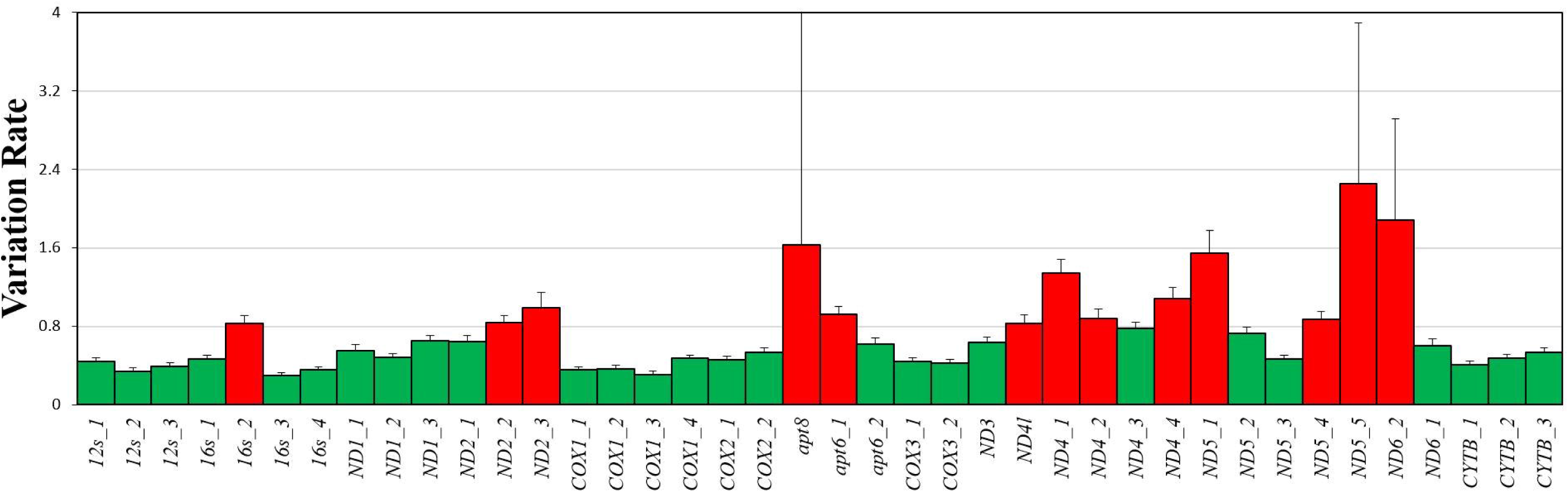
Variation rate cross-region in two rRNA and 13 protein-coding genes. The histogram represents variation rate across 40 loci in two rRNA and 13 protein-coding genes. Red column represent highly variable loci. The length of 16s_2 is 400 bp. The length from ND2_2 and ND2_3 is 699 bp. The length from apt8 and apt6_1 is 529 bp. The length from ND4l to ND4_2 is 996 bp. The length from ND4_4 to ND5_1 is 717 bp. The length from ND5_4 to ND6_2 is 1021 bp.

**Figure 2.**
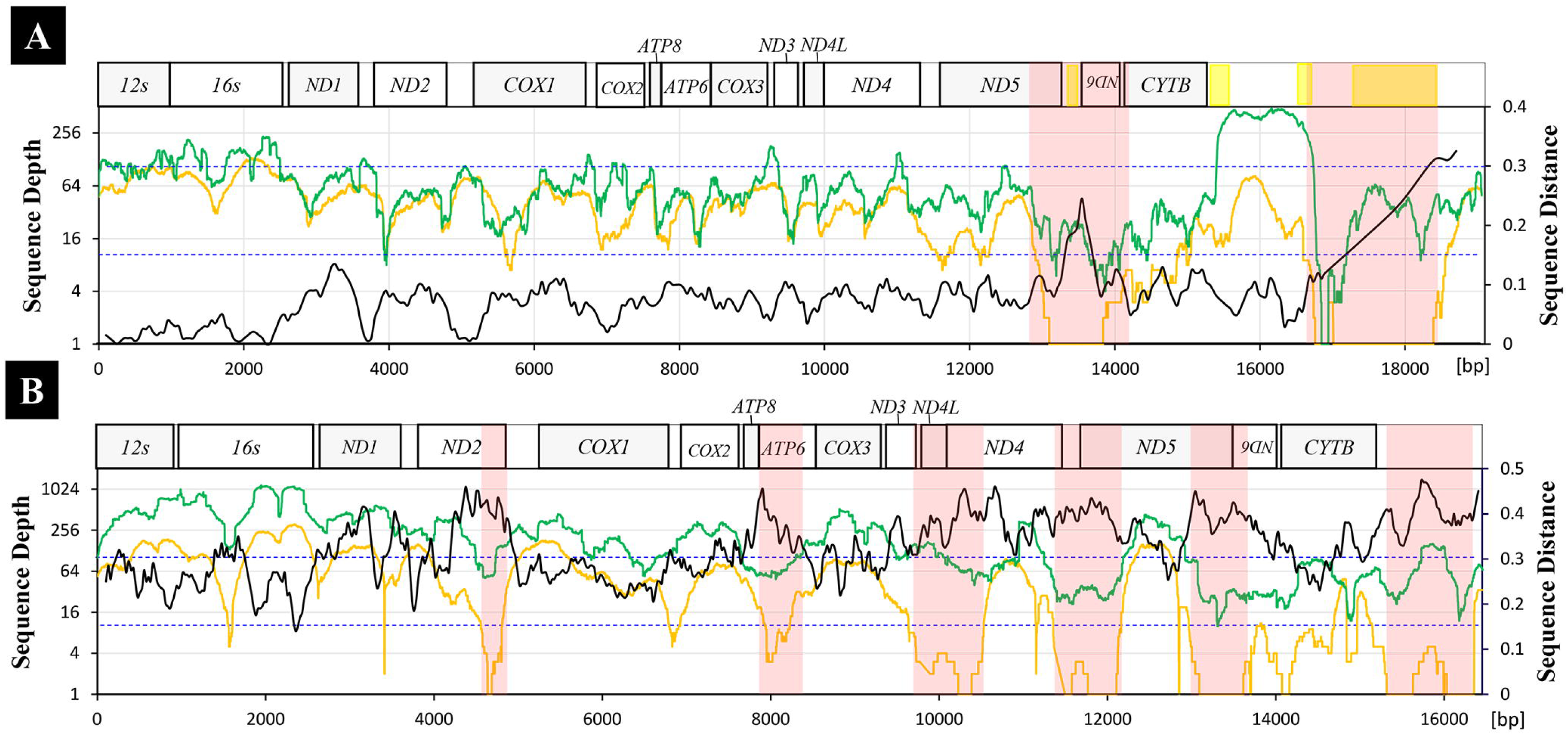
Coverage distributions for 400 bp and 2 kb library. **A** represents *Rana sp.1* results by using standard capture hybridization (orange line) and LR-HY (green line). Black line represents DNA sequence distance between *Rana sp.1* and *Rana sp.2* (Kimura 2-parameter distance). The sliding window length is 50 bp and the step length is 5 bp (below is the same). Dashed lines in A and B are constant at 0.15 and 0.3 sequence distance. The repetitive regions in *Rana sp2* which is labeled with yellow ranged from 13,424 to 13,572 bp, 15,402 to 15,660 bp, 16,593 to 16,770 bp and 17,382 to 18,498 bp. **B** represents *Onychodactylus sp.1* results by using standard capture hybridization (orange line) and LR-HY (green line). Black line represents DNA sequence distance between *Rana sp.1* and *Onychodactylus sp.1*. Dashed lines in A and B are constant at 0.15 and 0.3 of Kimura 2-parameter distance. The regions with greatest sequence-depth improvement are highlighted with red box.

**Figure 3.**
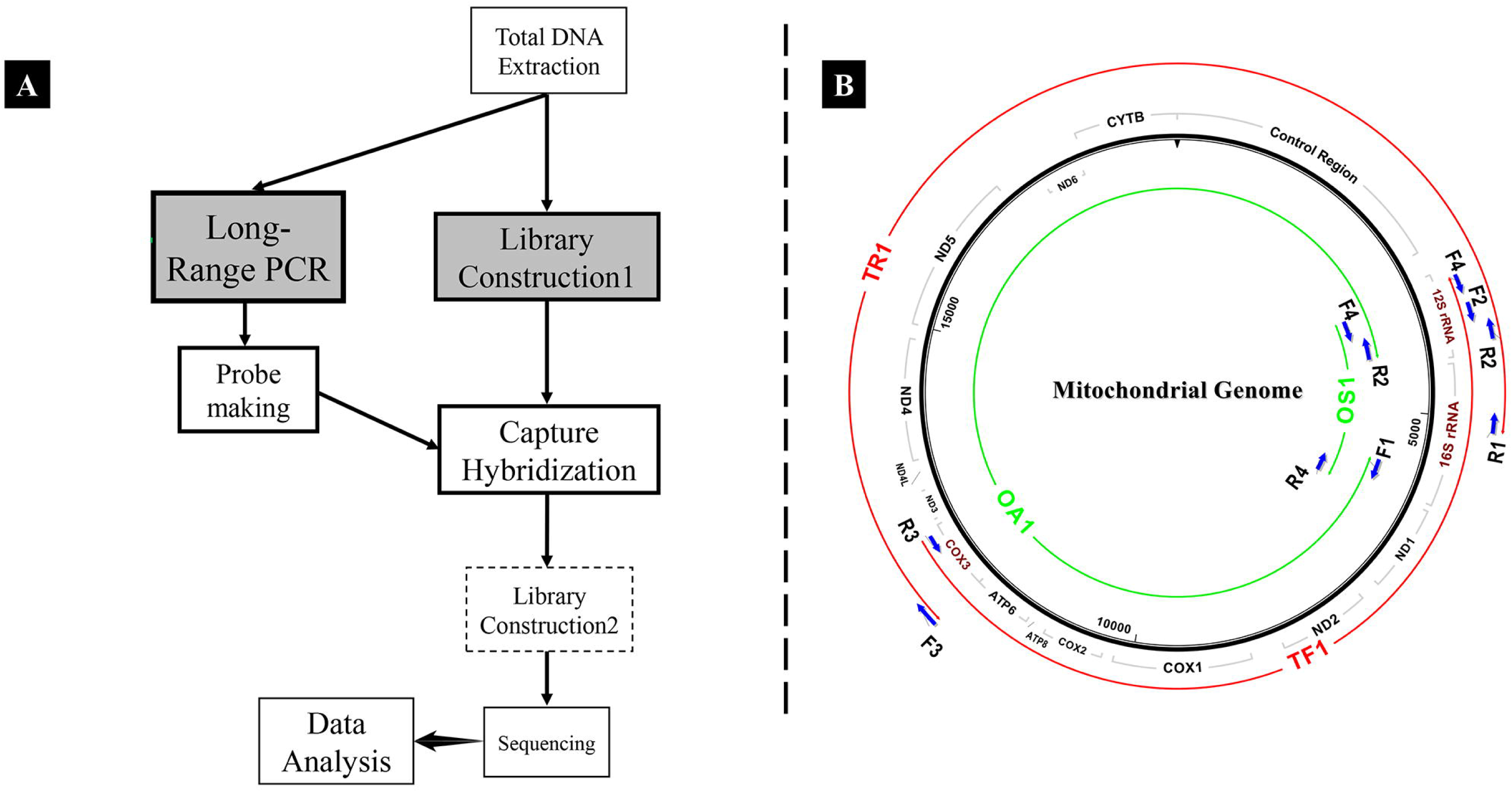
**A. Schematic pipeline for enriching mitochondrial DNA to high-throughput sequencing.** Compared to standard library, the long-range library hybridization strategy (LR-HY) has some modifications in library construction 1 and 2. LR-HY requires a long fragment during shearing in library construction 1 for capturing high variable loci and additional library construction 2 for Ion Torrent Personal Genome Machine sequencing. In standard hybridization, there is no construction library 2 and the enriched fragments directly sequenced; **B**. **Two strategies for amplifying mitochondrial genomes, termed OA1/OS1 and TF1/TR1**. OA1/OS1: amplification of OA1 and OS1 regions uses primers F1/R2 and F4/R4, respectively. TF1/TR1: amplification of fragments TF1 and TR1 using primers (F2/F4)/R3 and F3/(R1/R2), respectively

## 2. Materials

All reagents and plastic ware should be sterile.

1. LongAmp DNA Polymerase (New England BioLabs).
2. 2.5 mM dNTP (Takara).
3. WIZARD gel extraction kit (Promega).
4. Ampure Beads (Beckman).
5. Ion Xpress Barcode Adapter Kits from 1 to 96 (ThermoFisher).
6. Ion Plus Fragment Library Kit (ThermoFisher).
7. IonShear kit (ThermoFisher).
8. Agarose gel.
9. Human Cot-1 DNA (Agilent).
10. Hybridization buffer and blocking agent (from a Agilent aCGH kit).
11. Streptavidin beads (M-270, Invitrogen).
12. Tween-20.
13. 3M sodium acetate.
14. TE buffer (10 mM Tris, 1 mM EDTA, pH 8.0).
15. EBT and TET: 1× TE buffer, 0.05% Tween-20.
16. 1× bind and wash (BWT) buffer: 1 M NaCl, 10 mM Tris-Cl, 1 mM EDTA, 0.05% Tween-20, pH 8.0.
17. Hot wash (HW) buffer: 200 mL 10× PCR buffer, 200 mL MgCl 2 (25 mM), 1.6 mL H_2_O.
18. Library Amplification Kit (KAPA).
19. 2100 Bioanalyzer (Agilent).
20. Qubit 2.0 (Invitrogen).
21. 2% E-gel (Invitrogen).
22. Focused-ultrasonicator M220 (Covaris).
23. PCR reaction tubes.
24. Covaris microTUBE.
25. NanoDrop (ThermoFisher).
26. Magnetic rack.
27. Hybridization oven.
28. A thermal cycler.

## 3. Methods

### 3.1 Prepare Probe

1. Primes for amplifying mitogenome To achieve universality and avoid impact on the gene rearrangement, we designe degenerate primers on the conservative regions as shown in Table 1 (Design principle see Note 1). TF1 is amplified using (F2/F4)/R3 (expected length: 5–9 kb, figure 1B, red) and TR1 is amplified using F3/(R1/R2) (expected length: 7–12 kb, Figure 1B, red). Primer pair F1/R2 is used to amplify OA1 (expected length: >14 kb, Figure 1B, green), which covers all the protein-coding and control regions. Primer pair F2/R4 is used to amplify OS1 (expected length: >2 kb, figure 1B, green) covers two rRNA genes: *16s rRNA* and a portion of *12s rRNA*. (Note 2)
2. Long-range PCR Long-range PCR is conducted in 25 μL reactions and mix the following reagents:
  a. 0.8 μL forward primers (10 μM).
  b. 0.8 μL reverse primers (10 μM).
  c. 3 μL dNTP (2.5 mM).
  d. 1 μL LongAmp DNA Polymerase.
  e. 5 μL 5 × PCR buffer.
  f. 50–200 ng template.
  LR-PCR condition is as follows: initial incubate at 95 °C for 1 min, 30–32 cycles at 94 °C for 10 s, 58 °C for 40 s, and 65 °C extensive for variable times, a final extension at 65 °C for 10 min, and hold 10°C forever. Extension times are 3 min for OS1, 10 min for TF1 and TR1, and 16 min for OA1 (Figure 1B). Check PCR product by using 0.8% agorase gel.
3. Purify LR-PCR product by using WIZARD gel extraction kit.
4. Measure the concentration with a Nanodrop. The amounts of products should be up to 0.1-1.2 μg (To ensure enough quantity of PCR product see Note 3).
5. Mix PCR products according to amplicon length (and empirically adjusted according to sequence-depth). The ratio of the TF1 to TR1 amplicon is 5:8. The ratio of the OS1 to OA1 amplicon is 1:12. Probe making is conducted in 50 μL reactions and mix the following reagents:
  a. PCR product mixture 1.3 μg.
  b. 5 μL 10 × dNTP Mix.
  c. 5 μL 10 × Enzyme Mix.
6. Mix and centrifuge briefly (15,000 × g for 5 sec).
7. Incubate at 16 °C for 90 min (The time parameter setting is different from manufactory protocol, the reason in Note 4).
8. Add 5 μL Stop Buffer.
9. Add 1/10 volume 3 M sodium acetate and 2 volumes cold (−20 °C) ethanol to the reaction tube. Freeze at −70 °C for 30 min.
10. Centrifuge at 15,000 × g for 10 min. Carefully remove the supernatant with an pipettor and dry the pellet.
11. Resuspend the pellet in 50 μL H_2_O and precipitate the probe with sodium acetate and ethanol as described above.
12. Resuspend the probe in TE buffer and store at −20 °C.

**Table 1.**
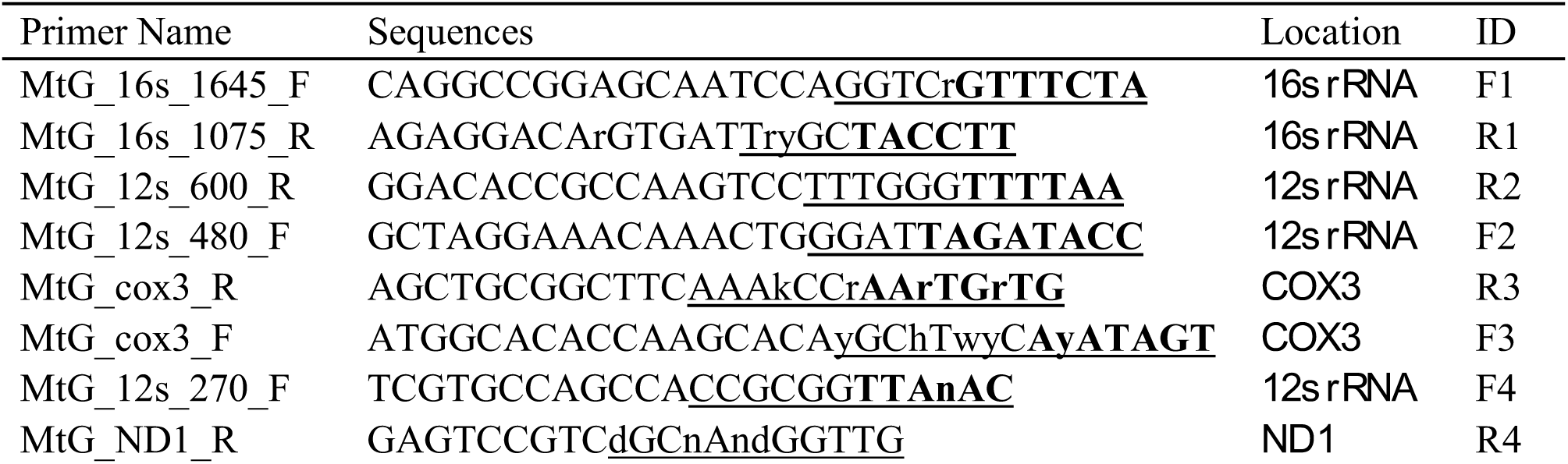
Primer information.

### 3.2 Library construction 1 for capture hybridization

1. LR-PCR products are mixed at a certain ratio the same with step 5 in the section Prepare Probe.
2. Shear the mixture in a Focused-ultrasonicator M220 (Covaris) by selecting the method DNA_2000bp_200_ul_Clear_microTUBE for 12 min. The shear volume is 200 μL (The shearing length can be adjusted and details is in Note 5).
3. End-repair reaction is carried out in 100 μL reactions and mix the following reagents:
  a. 130 ng sheared DNA.
  b. 20 μL 5 × End Repair Buffer.
  c. 1 μL End Repair Enzyme.
4. Adaptor ligation is carried out in 100 μL reactions and mix the following reagents in Ion Plus Fragment Library Kit:
  a. 130 ng of sheared DNA.
  b. 1.6 μL (Ion Xpress Barcode Adapter Kits from 1 to 96).
  c. 10 μL 10 × Ligase Buffer.
  d. 2 μL dNTP Mix.
  e. 2 μL DNA Ligase.
  f. 8 μL Nick Repair Polymerase. Incubate for 20 min at 25 °C in a thermal cycler and followed 72 °C incubation for 5 min.
5. Select long DNA fragment by using 0.4 volume of Ampure beads (i.e., 100 μL sample of DNA gets 40 μL of Ampure beads) (alternative choice is gel selection, see Note 6)
6. Library amplification is carried out in 100 μL reactions and mix the following reagents:
  a. Size-selected library;
  b. 10 μL 5 × PCR buffer;
  c. 5 μL 2.5 mM dNTP;
  d. 2 μL of 10 μM forward and reverse primers (Primer information in Note 7);
  e. 2 μL LongAmp DNA Polymerase. Incubate 95 °C for 1 min, then 15 cycles of 94 °C for 10 s, 58 °C for 40 s, 65 °C for 3 min, and finally 65 °C for 10 min followed by holding at 4 °C.
7. Purify with Ampure bead and add 15 μL × 1 TE buffer.

### 3.3 In-solution Capture hybridization

1. In-solution capture hybridization is carried out in 100 μL reactions:
  a. 25 μl 2 × hybridization buffer.
  b. 5 μl 10 × blocking agent.
  c. 2 μl Human Cot-1 DNA.
  d. 2 μl of blocking adaptors (from Ion Plus Fragment Library Kit, ThermoFisher).
  e. 10-100 ng of bait and 100-1000 ng library (Certain ratio of library and probe is 1:10). Incubate for 5 min at 95 °C, and then incubated for 72 hr at 65 °C.
2. After hybridization, incubate the mixture with 5 μL magnetic streptavidin beads (M-270, Invitrogen) for 20 min at room temperature.
3. Place the mixture into a magnetic rack to separate the magnetic beads from the supernatant.
4. Discard the supernatant.
5. Wash the beads using 200 μL of 1x BWT buffer, vortex the mixture for 30 s each time.
6. Discard the supernatant.
7. Repeat steps 5 and 6 for four times.
8. Wash the beads once with warmed HW buffer at 50 °C for 2 min.
9. Wash the beads once with 200 μL of 1x BWT buffer, vortex the mixture for 30 s.
10. Wash the beads once with 100 μL of TET, vortex the mixture for 30 s.
11. Separate hybridized target molecules from the bait in 30 μL TE by incubation at 95 °C for 5 min in a thermal cycler.
12. The PCR condition is the same with step 6 in section Library construction 1 for capture hybridization.

### 3.4 Library construction 2 for sequencing

1. Shear the enriched libraries for 120 s using an IonShear kit (ThermoFisher) in an open thermocycler. The 2 kb DNA fragments will be sheared to 300-500 bp.
2. Adaptor ligation is the same with step 4 in the section Library construction 1 for capture hybridization.
3. Select 450 bp reads by using 2% E-gel.
4. Library amplification is carried out in a PCR volume of 50 μL by using a Library Amplification Kit:
  a. 25 μL HiFi mix;
  b. 21 μL selected fragment solution;
  c. 4 μL primer mix (from Ion Plus Fragment Library Kit, ThermoFisher);
5. Concentration is measured by using Qubit. Length of library is measured by using 2100 Bioanalyzer (Agilent).

## 4. Notes

1. All of the primer structures are delicately refined. We separate the primers into two regions: the 5’ non-degenerate clamp region and the 3’ degenerate core region (11). The 3’ degenerate core region contains almost all the degenerate points for increasing the possibility mapped to the template. To stabilize the extension of the polymerase, we increase the GC content of the 5’ non-degenerate clamp and the AT content of at the beginning of the 3’ degenerate core region (12).
2. We recommend TF1 and TR1 amplification as *a prior* choice. If it failed, the OA1 also hard to success.
3. To assure the quantity of product enough for probe making, we recommend to amplify multiple tubes of LR-PCR instead of increasing cycles for a tube.
4. We tried short time of 20 min, 30 min, 40 min and 1h. Although the short time setting enable us to get long probe. But the efficiency of the probe is low. There is generally less than 10% reads mapped to reference genome.
5. Coveris provides many programs to shear DNA to different length from 50 bp to more than 10 kb. According the divergent gene length, we could adjust the shearing length. But it is not recommend to exceed >10 kb, because extremely high quality DNA samples are required. The long-range PCR also have high failure rate.
6. Using agarose gel to do size selection is another choice. But it loss more DNA and cost more time as compare to using Ampure beads.
7. Primers for Ion torrent library amplification can be booked by ourselves. Forward primer: 5’-CCATCTCATCCCTGCGTGTCTC-3’ and Reverse primer: 5’-CCACTACGCCTCCGCTTTCCTC-3’.

## Acknowledgments

This work was supported by the Ministry of Science and Technology of China (MOST no. 2012FY110800 to W.W.) and the National Natural Science Foundation of China (NSFC no. 31090251 to Y.Z.).

